# The bromodomain protein Bdf3 is sufficient to activate expression of a procyclin gene in bloodstream stage African trypanosomes

**DOI:** 10.64898/2026.01.12.698981

**Authors:** Evan J. Kim, Ethan L. Goroza, Ashley Y. Tan, Jolyne J. Lin, Kieran C. Saucedo, Fuminori Tanizawa, Jaclyn E. Smith, Danae Schulz

## Abstract

The early branching eukaryotic parasite, *Trypanosoma brucei*, lives extracellularly in the mammalian host, where it expresses variant surface glycoprotein (VSG). Antigenic variation of VSGs allows the parasite to evade the host immune response while in the bloodstream. Upon transition to the tsetse fly vector, the parasite remodels its surface to express procyclin protein from a limited set of *EP* and *GPEET* genes. While the environmental signals that trigger the switch from expressing *VSG* genes to expressing *EP* and *GPEET* genes are known, the molecular mechanism that initiates transcription of the *EP* and *GPEET* genes during parasite differentiation to the procyclic stage within the tsetse is not well understood. Previous work has shown that the chromatin interacting bromodomain protein Bdf3 is absent from the pol I promoter of the *EP* locus in bloodstream parasites, but appears there shortly after initiation of differentiation from the bloodstream to the procyclic stage. Here, we show that tethering of Bdf3 to the *EP* promoter using a dCas9-Bdf3 fusion protein and a guide RNA is sufficient to increase transcript levels of *EP1* in bloodstream parasites, where the gene is normally silenced. This result supports the model that Bdf3 appears at the *EP* locus during differentiation to facilitate initiation of transcription, which is consistent with the role for Bdf3 as a transcriptional activator in other systems. Understanding gene regulatory mechanisms in this early branching eukaryote may help build a more complete picture for the conserved and unique features of gene regulatory proteins across diverse biological systems.

**Importance:** The *Trypanosoma brucei* parasite is transmitted to a mammalian host via the tsetse fly vector, where it causes African trypanosomiasis, a fatal disease that causes a severe human and economic burden for people living in in areas of sub-Saharan African. While drug treatments have improved for some strains, some infections still require treatment with drugs that have serious side effects. One avenue for drug development is to try and manipulate the lifecycle of the parasite to make it poorly adapted to the mammalian host. However, this requires detailed knowledge of the mechanisms by which parasites regulate their genes as they progress through the lifecycle. Here, we show that the DNA interacting bromodomain protein Bdf3 may be a key driver for turning on insect-specific genes that code for insect-stage surface proteins. This finding could aid in the development of lifecycle manipulating drugs, and shed light on how gene regulatory mechanisms evolved.

## Introduction

Numerous parasites transition between hosts as p34r5tyart of their natural life cycle. Because host environments can differ dramatically with respect to temperature, pH, and nutrient availability, parasites that transition between hosts must evolve systems to adapt to differing environments. One such parasite is the African trypanosome, *Trypanosoma brucei*. *T. brucei spp.* are unicellular, eukaryotic protozoan parasites that infect humans and ungulates and cause Human and Animal African Trypanosomiasis disease. African trypanosomiasis is usually fatal, and imposes a severe human and economic burden for people living in Sub-Saharan Africa, where the disease is endemic (1, 2). *T. brucei* cycle between a tsetse fly vector and the bloodstream of a mammalian host, where they live extracellularly. Compared to the tsetse, the bloodstream environment of the mammalian host is warmer and glucose rich, and the parasite must contend with a robust antibody response mounted by the host immune system. The parasite has evolved to live in the mammalian bloodstream by antigenically varying the proteins on its surface, which are coded for by a large repertoire of Variant Surface Glycloprotein (*VSG*) genes (3). Antigenic variation of VSG surface proteins allows bloodstream parasites to successfully evade the host antibody response. Upon transition to the fly midgut following a tsetse bloodmeal, the parasites alter their glucose metabolism (4–7) and express a far more limited repertoire of procyclin surface proteins coded by the *GPEET* and *EP* genes (8). GPEET proteins contain a pentapeptide repeat of the GPEET amino acid sequence, while EP proteins contain a dipeptide repeat (8, 9). *GPEET, EP1, EP2,* and *EP3* are all co-expressed within the first few hours after differentiation to the procyclic form. GPEET protein predominates on the parasite surface early in development, and this protein is replaced with EP isoforms later in development within the midgut (10). Since the tsetse lacks an antibody-based immune response, antigenic variation of *VSG* genes is no longer required, and instead the GPEET and EP procyclin proteins shield the parasites from proteases expressed by the tsetse (11). Thus, remodeling of the parasite surface is essential to survive following transition to either the bloodstream or the tsetse environments.

While it has been established that expression of *GPEET* and *EP* genes is transcriptionally and developmentally regulated (12, 13), the molecular mechanism that initiates transcription of these genes following parasite transition to the fly is poorly understood. Unusually for a eukaryote, the *GPEET* and *EP* protein coding genes are transcribed by Pol I (12, 14–16). Because most protein coding genes in *T. brucei* are transcribed by Pol II in polycistronic units with limited transcriptional regulation (reviewed in (17) and recently corroborated with SLAM-seq (18)), the transcriptionally controlled procyclin genes offer a unique opportunity to examine transcriptional regulation in a highly diverged, early branching eukaryote.

Despite the lack of transcriptional control for pol II genes, *T. brucei* parasites do harbor much of the machinery associated with the histone code that is best characterized in model systems such as yeast, worms, and flies (19). This machinery includes numerous post-translational modifications of histone proteins (20) as well as proteins that read, write, and erase those modifications (21–28). One of the best characterized domains found in proteins that act as chromatin readers is the bromodomain, which recognizes acetylated lysines on histone tails and has an established role in gene activation (29–33). In murine models, bromodomain proteins maintain cell identity and help regulate differentiation (34–36).

Bromodomain proteins have also been examined in protozoan parasites, where it has been found that inhibition via small molecules or genetic manipulation results in defects in transcription for *Toxoplasma gondii* (37) and *Leishmania mexicana* (*38*) as well as developmental blocks in *Trypanosoma cruzi* (39, 40) and *Plasmodium falciparum* (41, 42). Small molecule inhibition of bromodomain proteins causes defects in egg development in *Schistosoma japonicum* (*43*). In *T. brucei*, bromodomain proteins are also involved in differentiation processes, both in bloodstream (25) and in procyclic (44) stages. *T. brucei* contains 7 proteins with bromodomains identified through homology (21, 26, 45). Bdf1-6 bind transcription start sites in bloodstream parasites (21, 25, 26), while Bdf7 localizes to transcription termination regions (26). Staneva et al. classified Bdf1, Bdf3, and Bdf4 as Class I transcription start region associated factors because of the sharp, high peaks observed in ChIP-seq experiments using antibodies against Bdf1, Bdf3, and Bdf4. Based on their genomic distribution, the authors hypothesized that Bdf1, Bdf3, and Bdf4 were likely to be involved in transcription initiation (26). Bdf2, Bdf5, and Bdf6 showed broader distributions, and thus were classified as Class II transcription start region associated factors. Bdf2, Bdf5, and Bdf6 were hypothesized to be involved in transcriptional elongation based on this distribution (26). Based on their genomic localization at pol II transcription start sites and association with transcriptional activation, we hypothesized that bromodomain proteins in *T. brucei* might also be important for initiating transcription of pol I driven procyclin genes that are transcriptionally activated as parasites transition from the bloodstream to the insect stage.

Using CUT&RUN (46), we followed the genomic localization of the bromodomain protein Bdf3 in pleomorphic parasites induced to differentiate from the bloodstream to the procyclic stage via treatment with cis-aconitate and incubation at 27°C (47). We found that both the *EP* locus and the *GPEET* locus lacked genomic occupancy of Bdf3 in bloodstream parasites. 3 hours after differentiation was induced, we observed *de novo* occupancy of Bdf3 at both the *EP* locus and the *GPEET* locus. Indeed, these were the only loci analyzed where Bdf3 was consistently absent from a genomic site in bloodstream parasites but present following induction of differentiation (48). While the presence of Bdf3 at the *EP* and *GPEET* loci implicates the protein as contributing to initiation of transcription at that locus, its mere presence does not definitively prove its ability to increase transcript levels of procyclin genes. We thus set out to test whether Bdf3 was sufficient to increase transcript levels at the *EP1* locus in bloodstream parasites, where the locus should normally be silenced. We engineered a tethering system inspired by CRISPR activation-based systems developed for other organisms (49, 50) and introduced it into a previously validated reporter system for *EP1* transcript levels (51). This system allowed us to artificially drag Bdf3 to the *EP1* locus in bloodstream parasites, where *EP1* is normally silenced. Here, we show that Bdf3 is sufficient to increase transcript levels of *EP1* under these conditions. This system can be used as a platform to study transcriptional activation for a host of other putative activators.

## Results

### The dCas9-Bdf3 fusion protein is inducibly expressed in *Trypanosoma brucei*

Previous work in our lab showed that Bdf3 is absent from the *EP1* promoter in bloodstream parasites (48). After treatment with cis-aconitate and incubation at 27°C to induce differentiation to the procyclic form, Bdf3 localizes to the *EP* locus at 3 hours post induction and remains there until differentiation is completed at 72h post-induction (48). However, whether Bdf3 occupancy at the *EP1* locus is sufficient to increase transcript levels of the *EP1* gene is unknown. In order to test whether Bdf3 is sufficient to induce expression of *EP1* in bloodstream parasites, we cloned a construct that would allow us to artificially recruit Bdf3 to the *EP1* locus in *EP1/GFP* reporter bloodstream parasites (51). To do this, we fused the entire Bdf3 open reading frame to a catalytically dead version of the Cas9 protein (dCas9) using a flexible GS linker (52). The dCas9 protein is recruited to a specific genomic locus using a guide RNA, but two mutations in the active site prevent dCas9 from creating a double strand break at that location (49, 53). In model systems, fusion of dCas9 to transcriptional activation domains (CRISPR activation) or transcriptional inhibition domains (CRISPR inhibition) and targeting of these fusion proteins to a gene promoter using a guide RNA modulates transcription for the target gene (49, 50). CRISPR activation and CRISPR inhbition systems have been successfully adapted in mammalian cells, plants (54), and in the protozoan parasite *Plasmodium falciparum* (55). Excellent work adapting CRISPR-Cas9 systems for genome editing in *T. brucei* (56–58) allowed us to take advantage of previously built constructs to set up the CRISPR activation system. To our knowledge, we are the first to adapt a dCas9-based CRISPR-activation system to *T. brucei*.

To address whether Bdf3 can increase transcript levels at a silent locus, our FLAG tagged dCas9-Bdf3 fusion construct was cloned into a vector that allows inducible expression using the tetracycline-on system (tet-on) (Fig. 1A, top) (59). The construct was stably integrated into the genome using homology to the rDNA locus. An additional FLAG tagged dCas9 construct was integrated to use as a control. Treatment for 1 day or 2 days with doxycycline (dox) showed inducible expression of both the dCas9-Bdf3 fusion protein and the dCas9 protein in transfected parasites, with both proteins appearing at the expected size (Fig.1B, Fig. 1C). We did not observe background expression of dCas9-Bdf3 in the - Dox condition for any of the three clones (Fig. 1B). We observed a robust expression increase in dCas9 protein levels after doxycycline treatment, although a faint band was observed in both clones in uninduced samples (-dox). Thus, we proceeded with clone 1. As an additional control, we cloned the Bdf3 open reading frame into the same vector that was used for both dCas9-Bdf3 and dCas9 alone and tested inducible expression by western blot (Fig. 1D). We observed the lowest level of background expression in Clone 2, as evidenced by undetectable protein in the -Dox condition, and so further experiments continued with this clone. Once inducible expression was verified for both the dCas9-Bdf3 strain and the dCas9-alone strain, we transfected both strains with guide RNAs targeted to either the *EP1* promoter or a locus unrelated to differentiation. We chose the *NEO* locus to use as the unrelated control. The *NEO* gene was previously used to select for integration of the T7 polymerase and tet repressor (59), which is present in the background of the *EP1/GFP* reporter strain. Guide RNA constructs were designed to target the *EP1* promoter (12, 14, 16, 60) or the *NEO* gene using the Eukaryotic Pathogen CRISPR guide RNA/DNA Design Tool (61). Tet/dox inducible guide RNA constructs were stably integrated into the trypanosome genome using selection with phleomycin (Fig. 1A), as in (56).

**Figure 1.**
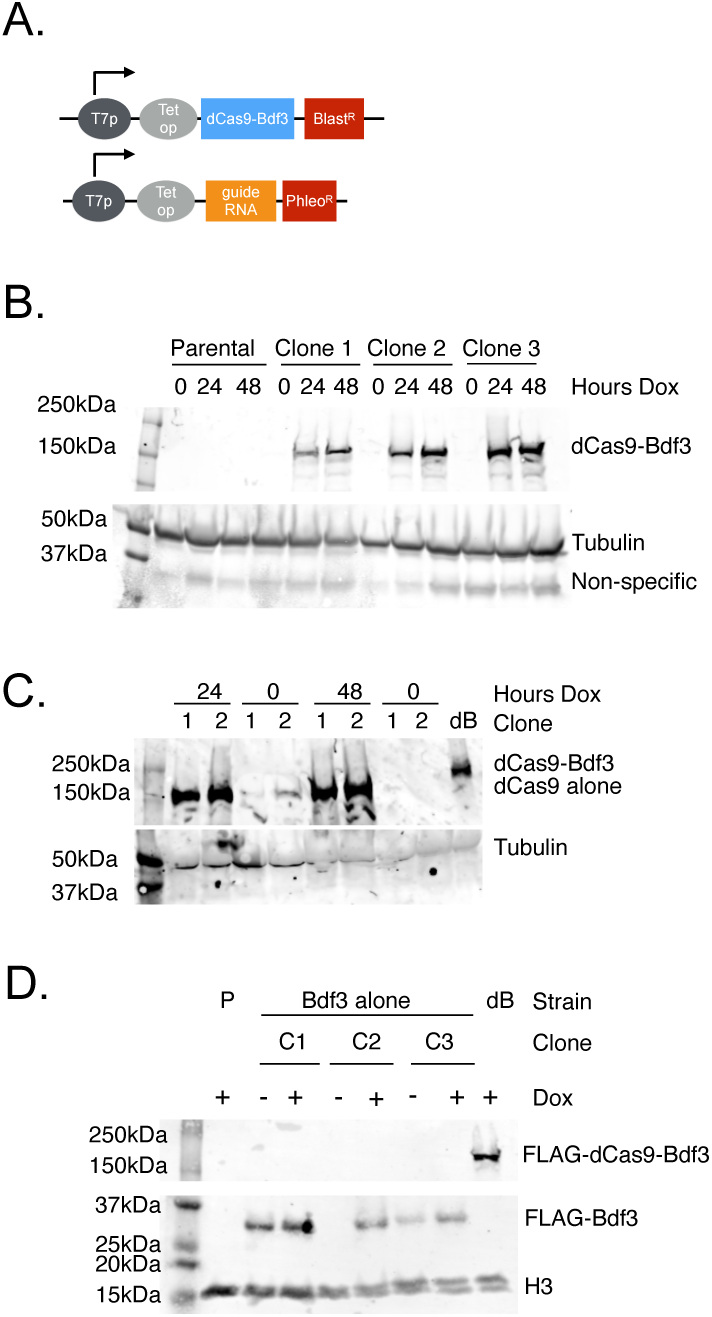
The dCas9-Bdf3 fusion protein and the dCas9 protein are inducibly expressed. (A) Schematic of constructs integrated into the *T. brucei* genome in order to inducibly express the dCas9-Bdf3 fusion protein and the guide RNAs. T7p, T7 promoter, Tet op, tetracycline operator Blast^R^, *BSD* gene to generate resistance to blasticidin. Phleo^R^, *BLE* gene encoding resistance to phleomycin. Both constructs are targeted to the rDNA locus. (B) Western blot of *T. brucei* extracts transfected with the dCas9-Bdf3 fusion construct and harvested at the indicated timepoints following treatment with 1µg/ml dox. 3 independent clones were analyzed along with a parental control that was not transfected with the dCas9-Bdf3 construct (labeled Parental). Blots were probed with anti-FLAG to detect dCas9-Bdf3 and anti-tubulin to detect tubulin protein. (C) Western blot of *T. brucei* extracts transfected with the dCas9 alone construct and harvested at the indicated timepoints following treatment with 1µg/ml dox. 2 independent clones were analyzed and an extract containing a dCas9-Bdf3 construct was used as a positive control (labeled dB). Blots were probed with anti-FLAG to detect dCas9 and dCas9-Bdf3 and anti-tubulin to detect tubulin protein as a loading control. (D) Western blot of *T. brucei* extracts transfected with the Bdf3 construct and harvested after 48h of treatment with 1µg/ml dox. – indicates untreated samples. 3 independent clones were analyzed (labeled C1, C2, and C3) along with a parental control that was not transfected with the Bdf3 construct. An extract containing a dCas9-Bdf3 construct was used as a positive control (labeled dB). Blots were probed with anti-FLAG to detect Bdf3 and dCas9-Bdf3 and anti-H3 to detect histone H3 as a loading control.

### The bromodomain protein Bdf3 is sufficient to increase transcript levels of *EP1/GFP* when targeted to the *EP1* locus

To ascertain whether Bdf3 is sufficient to increase transcript levels of *EP1/GFP* bloodstream parasites where it should not be highly expressed, we measured *EP1/GFP* levels by flow cytometry in *EP1/GFP* reporter parasites harboring either *EP1* guides or *NEO* guides in both dCas9-Bdf3 strains and dCas9 alone strains. 3 different clones were chosen to analyze for each guide RNA tested, and dox was used to induce expression of both the guide RNA and either the dCas9-Bdf3 fusion protein, or the dCas9 alone protein. Flow cytometry measurements were taken after 48h and 72h of dox treatment. For most clones, additional biological replicates were analyzed, except in the case where a clone was inadvertently lost. For each sample, the mean GFP fluorescence intensity was obtained for the +Dox treatment where dCas9-Bdf3, dCas9 alone and the guide RNA were induced, and also the uninduced -Dox treatment used as the control. The fold change in mean GFP fluorescence intensity for the +Dox versus the -Dox samples was computed and is shown in Fig. 2. We found that there was a significant increase in +Dox/-Dox fold changes for mean GFP fluorescence intensity at both time points in dCas9-Bdf3 parasites targeted to the *EP1* locus using two different guide RNAs (dCas9-Bdf3 EP1-1 and dCas9-Bdf3 EP1-3) when compared to control dCas9-Bdf3 parasites targeted to the *NEO* locus (dCas9-Bdf3 NEO-1) (Fig. 2, S1 Fig., S1 Table, S2 Table). This result indicates that the increase in *EP1/GFP* expression occurs only when the fusion protein is targeted to the *EP1* locus, and is thus locus dependent. We also found that there was a significant increase in +Dox/-Dox fold changes in mean GFP fluorescence intensity when dCas9-Bdf3 parasites targeted to the *EP1* locus were compared to dCas9 alone parasites targeted to the *EP1* locus at both timepoints, indicating that dCas9 alone is not sufficient to increase *EP1/GFP* and that Bdf3 is also required (Fig. 2, S1 Table, S2 Table). We do not have evidence that overexpression of Bdf3 alone is sufficient to increase transcript levels of *EP1/GFP*, as there was no significant difference in mean GFP fluorescence intensity fold changes for Bdf3 alone parasites compared to dCas9 alone parasites targeted to either *NEO1* or *EP1* (S1 and S2 Tables). Lastly, there is no significant difference in +Dox/-Dox fold changes in dCas9 alone parasites targeted to *EP1* compared to dCas9 alone parasites targeted to the *NEO* locus. We conclude that Bdf3 is sufficient to increase transcript levels of *EP1/GFP* when targeted to the *EP1* locus. Given that Bdf3 appears at the *EP1* locus after differentiation from the bloodstream to the procyclic stage is initiated (48), our data support a model in which Bdf3 facilitates transcription at the *EP1* locus once differentiation has been triggered.

**Figure 2.**
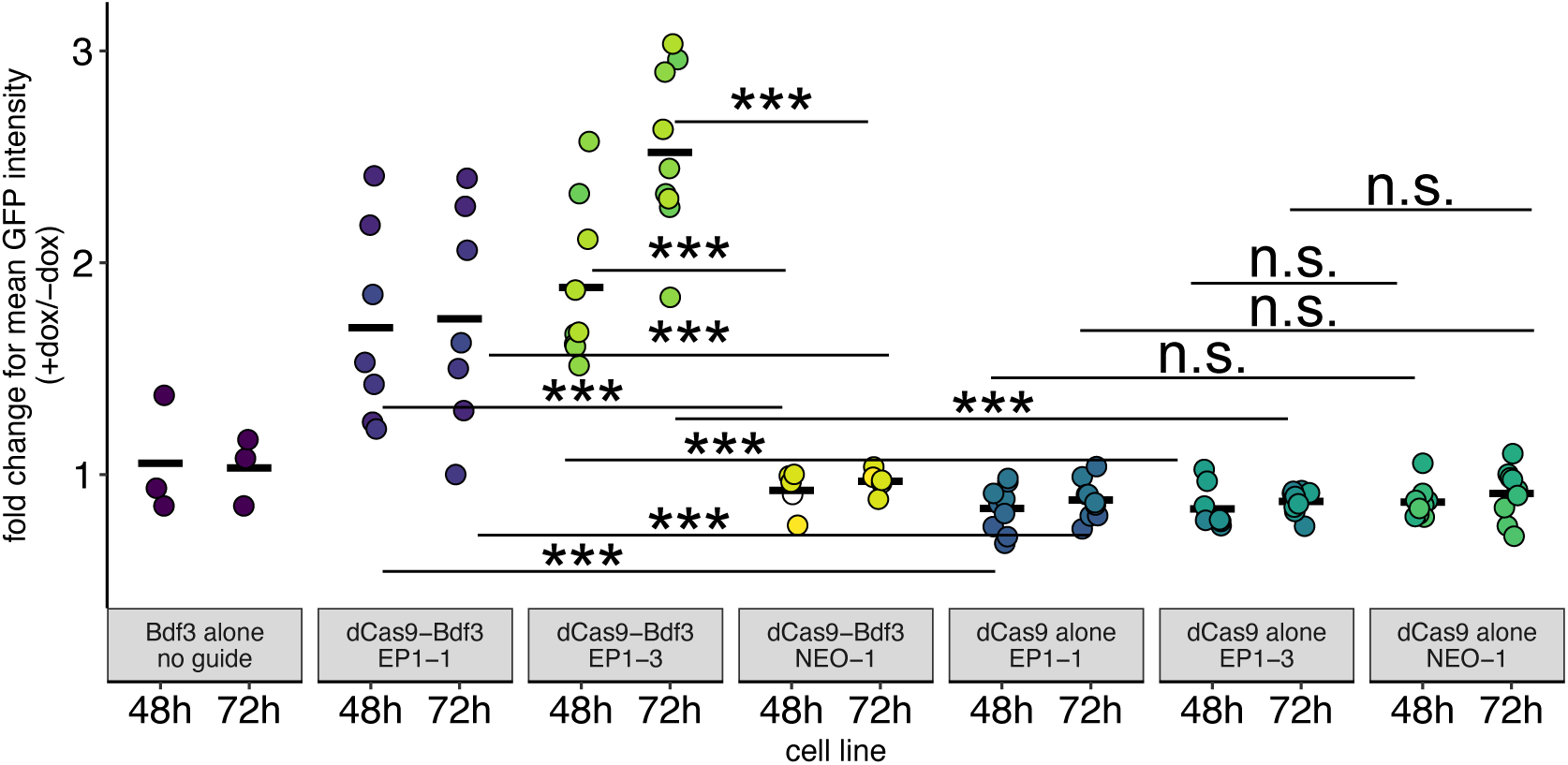
dCas9-Bdf3 targeted to the *EP1* locus is sufficient to increase *EP1/GFP* transcript levels in bloodstream *T. brucei* parasites. Quantification of flow cytometry data processed to compute the fold change in mean fluorescence intensity of parasites treated with dox vs untreated parasites. 1µg/ml dox treatment was used to induce expression of dCas9-Bdf3, dCas9 alone, Bdf3 alone and guide RNAs targeted to either the *EP1* locus (EP1-1 and EP1-3 guides) or the *NEO* locus (NEO-1 guide). 3 independent clones were analyzed for each the dCas9-Bdf3 and dCas9 alone strains. Only one clone of Bdf3 alone was analyzed with 3 biological replicates. Dots are color coded such that biological replicates for the same clone are the same color. Data was analyzed using ANOVA followed by a Tukey HSD (Honestly Significant Difference) post-hoc analysis. Two different ANOVA analyses were performed, one for 48h and one for 72h dox treatment. *, p < 0.05, **, p < 0.01, ***, p < 0.001, n.s. not significant. The figure indicates the most relevant statistical comparisons. A full set of p values for each comparison is provided in S1 Table and S2 Table.

### Expression of the dCas9-Bdf3 fusion protein targeted to the *EP1* locus causes a mild growth defect

In order to assess whether expression of *EP1/GFP* induced by dCas9-Bdf3 causes a growth defect, we performed growth assays on both *EP1* and *NEO* guide RNA parasites expressing either the dCas9-Bdf3 fusion protein or dCas9 alone. We observed a mild growth defect in dCas9-Bdf3 parasites targeted to *EP1* after 48h of induction, while this growth defect was not observed in any other strain we tested (Fig. 3). We do not think the growth defect is caused solely from the expression of the dCas9-Bdf3 fusion protein because parasites induced to express dCas9-Bdf3 targeted to the *NEO* locus do not exhibit a growth defect.

**Figure 3.**
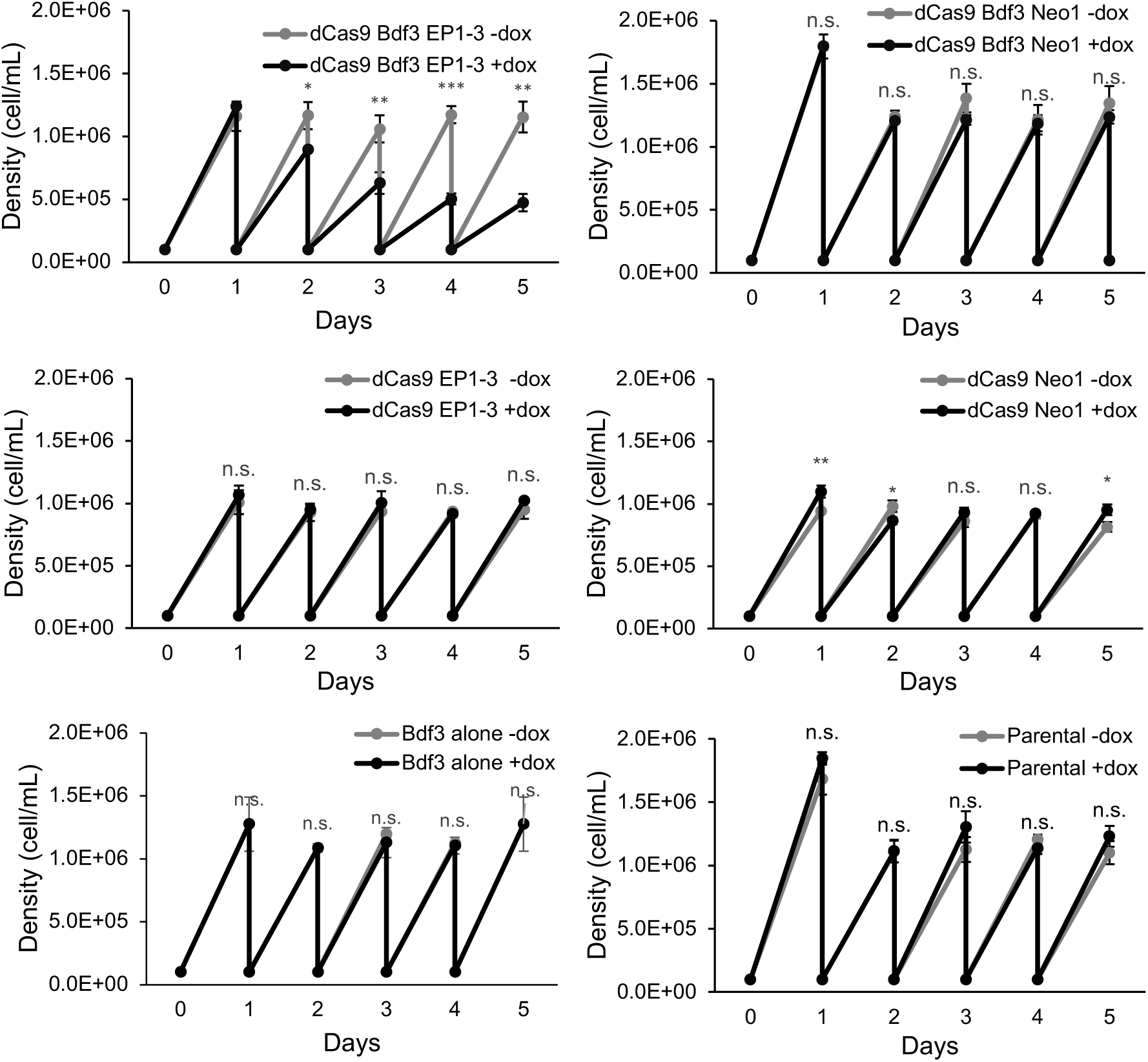
Expression of dCas9-Bdf3 targeted to the *EP1* locus causes a growth defect after induction of dCas9-Bdf3 and the guide RNA. Parasites were seeded at 100,000 cells/ml with or without 1µg/ml dox and grown for 24h before being counted and diluted back to 100,000 cells/ml over the course of 5 days. A Student t-test was used to compared -dox and + dox samples for each parasite strain at each time point. *, p < 0.05, **, p < 0.01, ***, p < 0.001. n.s. not significant.

Multiple researchers have shown that expression of more than one VSG protein can lead to growth defects (62–65), and it’s possible that the expression of both *VSG* and *EP1* might cause a similar phenotype. Another possibility relates to the growth arrested nature of stumpy parasites, which are an intermediate stage that occurs when bloodstream parasites grow to high density; these parasites have been shown to pre-express *EP1* prior to transition to the fly (66). While the monomorph strains we used in this experiment do not form a true stumpy intermediate, the increased transcription of *EP1* might cause the parasites to acquire some stumpy-characteristics, causing a reduction in growth. Small molecules that induce expression of *EP1* have also been associated with growth defects (51). A study using a procyclin reporter also found that some compounds that induced procyclin reporter activity also inhibited growth of the parasites (67). Thus, while we do not know the exact mechanism for the observed growth defect in dCas9-Bdf3 parasites targeted to the *EP1* locus, other studies have shown that growth inhibition is often associated with an increase in *EP1* transcript levels.

### Transcript levels of *EP1, GPEET, and PAG2* are increased after expression of dCas9-Bdf3 targeted to the *EP1* locus

To verify the flow cytometry results, we measured *EP1* transcript levels directly using q-PCR on parasites induced to express dCas9-Bdf3 targeted to the *EP1* locus or to the control *NEO* locus following 3 days of Dox treatment. We found that both *EP1* and *GFP* transcript levels were higher in parasites induced to express dCas9-Bdf3 and a guide RNA targeted to *EP1* compared to uninduced parasites (Fig. 4A). Parasites induced to express dCas9-Bdf3 and a guide RNA targeted to the control *NEO* locus did not show an increase in expression of either *EP1* or *GFP*, as expected (Fig. 4B). In addition, expression of a control gene *SF1* (splicing factor 1) that was not expected to change following induction of dCas9-Bdf3 showed no significant change in expression when dCas9-Bdf3 was targeted to either the *EP1* locus or the *NEO* locus (Fig. 4A, Fig. 4B). Because the promoter sequence for *EP1* and *GPEET* are highly similar, we expected that guide RNAs targeted to the *EP1* promoter may also tether dCas9-Bdf3 to the *GPEET* locus, resulting in an increase in *GPEET* expression when both dCas9-Bdf3 and the guide RNA are induced with dox. This hypothesis was supported by an increase in expression of the *GPEET2* gene following induction of both dCas9-Bdf3 and the *EP1* guide RNA (Fig. 4A). No increase in *GPEET2* expression levels was observed in parasites induced to target dCas9-Bdf3 to the *NEO* locus. Given that the *PAG2* gene is found in the same polycistronic unit as *EP1*, we measured the expression levels of *PAG2* following induction of dCas9-Bdf3 targeted to the *EP1* locus. Expression levels of *PAG2* were significantly increased in dCas9-Bdf3 parasites expressing guide RNAs targeted to *EP1* compared to uninduced parasites (Fig. 4A). These changes were not observed when dCas9-Bdf3 was targeted to the control *NEO* locus (Fig. 4B). The observed increase in *EP* and *GPEET* expression in parasites expressing dCas9-Bdf3 targeted to the *EP1* locus are not as large as the changes observed in expression of these genes as parasites are fully differentiating from the bloodstream to the insect stage (51, 68, 69). This observation is consistent with the fact that multiple modes of stabilizing *EP* and *GPEET* transcripts are in play in fully differentiating parasites. Stabilization of *EP* and *GPEET* transcripts has been shown to be mediated through UTR sequences (70–72) and trans-acting factors (73, 74). Environmental factors such as nutrient availability can also influence *GPEET* transcript stability (72, 75). Thus, it is not surprising that transcript levels of *GPEET* and *EP1* observed in dCas9-Bdf3 *EP1* targeted parasites are not as high transcript levels observed in differentiating parasites. Finally, multiple different bromodomain protein complexes have been shown to localize to pol II promoters (26). It is possible that additional transcriptional activators, in addition to Bdf3 and its associated complex members, are required to achieve high transcript levels at the *EP1* promoter or pol I promoters in general.

**Figure 4.**
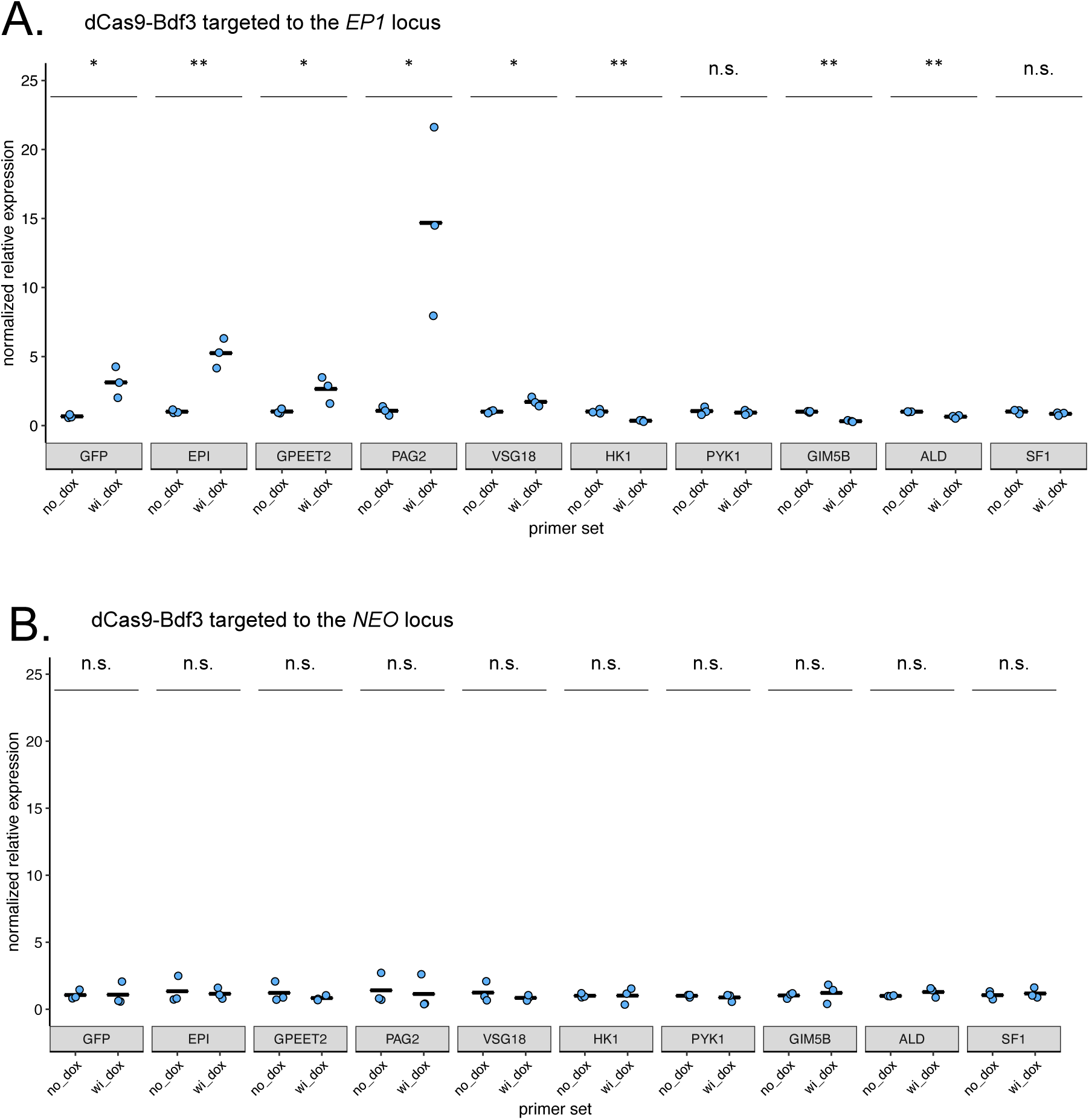
dCas9-Bdf3 targeted to the *EP1* locus results in increased expression of *EP-*related genes. Relative expression of each indicated gene analyzed using quantitative PCR from cDNA generated from RNA extracts of parasites harboring dCas9-Bdf3 targeted to the *EP1* locus (A) or dCas9-Bdf3 targeted to the *NEO* locus (B). Parasites were treated with dox for 72h to induce maximal expression of dCas9-Bdf3 and the guide RNA. Scaling was performed such that the average -Dox value for each gene was set to 1 to allow for easy comparisons between samples. Each dot represents the average of 3 technical replicates and the set of 3 dots for each sample represents 3 biological replicates for 3 independent clones. A Student t-test was performed to compare +Dox samples vs -Dox samples. *, p < 0.05, **, p < 0.01, ***, p < 0.001, n.s. not significant. wi_dox, with dox

Previous work in our lab has showed that treatment of *T. brucei* parasites with either spironolactone or eflornithine causes an increase in expression of *EP1, GFP, and PAG2,* and that these changes are accompanied by other changes in transcript levels that occur as parasites transition from the bloodstream to the insect stage (51). These changes include a decrease in expression of genes that are important in the glycolysis pathway, including *PYK1, HK1, ALD*, and *GIM5B*. We were surprised to see that transcript levels of *HK1, GIM5B and ALD* were slightly decreased in parasites where dCas9-Bdf3 was induced and targeted to the *EP1* locus (Fig. 4A). No significant changes in transcript levels for these genes was observed in parasites where dCas9-Bdf3 was induced and targeted to the control *NEO* locus (Fig. 4B). A silent *VSG* gene (VSG18) was also slightly upregulated, which has also been previously observed in conjunction with changes in *EP1* expression (25). The unexpected decrease in transcript levels for genes associated with glycolysis may indicate that a feedback loop of unknown mechanism may exist in parasites for which *EP1* transcript levels are increased. Increases in transcript levels for *EP1* and *GPEET* have been observed in conjunction with a decrease in transcript levels for glycolysis genes in stumpy parasites that undergo transcriptional pre-adaptation to life in the tsetse (66, 68), so there is precedent for these changes occurring together. However, we did not necessarily expect a differentiation program to initiate following an increase in *EP1* transcript levels, and this phenomenon might be worth exploring in future work. It should also be emphasized that the observed difference in transcript levels for glycolysis genes is quite muted compared to the much more dramatic changes in transcript levels observed in bloodstream parasites induced to differentiate to the procyclic form (68, 69).

## Discussion

While the role for bromodomain proteins as transcriptional activators in mammalian cells is well established, whether these proteins are important for facilitating transcription initiation in trypanosomes is less clear. Here, we show that the bromodomain protein Bdf3 is sufficient to increase transcript levels for a gene that is normally silenced in bloodstream parasites (Fig. 2). Both Bdf3 and a guide RNA targeting the *EP1* promoter are required to observe an increase in *EP1* transcript levels (Fig. 2).

One thing that remains unclear is whether acetylation at the promoter of the *EP1* locus is required for Bdf3 to increase transcript levels. In many model systems, bromodomains and histone acetylation domains are often found in one protein, allowing both acetylation of histones and reading of that acetylation to be accomplished by the same protein. Examples include p300, CBP, PCAF, and GCN5, among others (76). In *T. brucei*, none of the 7 identified proteins with bromodomains also contain a HAT domain, but there are 3 other identified *T. brucei* proteins with HAT domains (22). Elegant experiments using immunoprecipitation combined with mass spectrometry have shown that Bdf3 complexes with both Bdf5 and Hat2 (26). Depletion of Hat2 in *T. brucei* alters the site of transcription initiation in *T. brucei* (77). Hat2 has been shown to acetylate H4K10 (22, 77), which is found at pol II promoters (21), and recruitment of Hat2 by Bdf3 might result in this same acetylation at the *EP1* promoter in our experiments. Bdf3 has also been shown to be important for deposition of H2A.Z at transcription start sites for pol II promoters (21, 78). However to our knowledge, aside from Bdf3 (48), it remains unclear whether the histone modifications and proteins observed at pol II promoters are also found at the pol I promoters of *EP* and *GPEET* as parasites differentiate from the bloodstream stage to stumpy forms and then to procyclic stages. This is an interesting area for future study. Our favored model is that increases in H4K10ac at the *EP1* promoter may drive recruitment of Bdf3 that is observed in differentiating parasites. Resynthesis of an H4K10ac-specific antibody would provide the tools to interrogate this question.

To our knowledge, we are the first group to engineer a CRISPR activation-type system in *T. brucei*. It remains unclear whether the tool could be easily adapted to overexpress proteins of interest. Because most pol II genes are expressed polycistronicly from transcription start sites, designing a guide to a specific transcription start site and increasing transcription at that site would be expected to increase transcription of multiple genes simultaneously. In addition, because most transcript levels are regulated post-transcriptionally through 5’ and 3’ UTRs (79, 80), pseudouridination (81), lncRNAs (82), and alternative polyadenylation (83), increasing transcription at the relevant start site may not have any effect on transcript levels for the gene of interest if these other modes of regulation dominate. If a construct containing the gene of interest is placed downstream of a unique promoter and incorporated exogenously, overexpression may be achievable through expression of the dCas9-Bdf3 fusion in conjunction with a guide RNA targeted to the promoter. However, the already existing tet inducible system (59) may offer the same advantages with less engineering.

One outstanding question that should be addressed in future work is whether Bdf3 is capable of inducing expression from any promoter, or whether there is something unique about the *EP1* locus. This issue has been difficult to address because only a very limited number of genes are transcriptionally regulated in different life cycle stages. In order to rigorously address this question, the promoter in question would ideally be silent in at least one life cycle stage. The promoters upstream of silent *VSG* genes are one example of a region that is silenced in bloodstream parasites, but because the expression site promoters share a great deal of homology, trying to target dCas9-Bdf3 to these promoters may be difficult because the protein would be diluted among many different sites. Another way to address this could be to design a guide RNA to an alternate exogenously introduced promoter. Many *T. brucei* expression constructs utilize either the *GPEET* or the *EP* promoter to drive expression, but utilization of the rRNA promoter is also common. Designing guide RNAs targeting the rRNA promoter and examining the effect on transcript levels of the adjacent gene would address whether Bdf3 is sufficient to increase transcript levels at multiple promoter types. However, because the rRNA promoter is strong and constitutively active in bloodstream parasites, it is possible that the addition of dCas9-Bdf3 would not boost the signal significantly if it is already ‘maxed out’, so to speak. Thus, a negative result for this experiment may be difficult to interpret. Because *NEO* is controlled by readthrough transcription in our trypanosome system, we did not attempt to control *NEO* expression using guide RNAs. Guide RNAs targeted to the *NEO* gene body were simply used as a sink for dCas9-Bdf3 to avoid untargeted interactions with other regions of the genome, something that is a known issue in the absence of a decoy or irrelevant guide RNA (84, 85).

Another useful feature of the engineered dCas9-Bdf3 system is that mutation analyses can now be used to determine what features of the protein are necessary to achieve the increased transcript levels observed when Bdf3 is targeted to the *EP1* locus. Mutation of a conserved tyrosine and asparagine (86, 87) in the bromodomain would allow interrogation for the requirement to bind acetylated histones. Truncations might also assist in determining what portions of the protein are necessary to recruit Hat2 and Bdf5, if any. Finally, Bdf3 is one of six trypanosome bromodomain proteins shown to localize to transcription start sites (21, 25, 26). The dCas9 fusion system that we have engineered can be used to ascertain whether other bromodomain proteins are sufficient to increase transcript levels at the *EP1* locus. The system can also be used to help characterize the many chromatin associated proteins found in complexes that localize to transcription start sites (26).

## Materials and Methods

### Strain details and culture growth

Bloodstream SM *EP1/GFP* parasites (51) were cultured in HMI-9 (88) at 37℃ and 5% CO_2_. The dCas9-Bdf3 fusion construct, the dCas9 alone construct, and the Bdf3 alone constructs were transfected into reporter parasites using an AMAXA nucleofector with the X-001 setting. For each transfection, 30-50 million parasites were harvested and resuspended in 100 uL of nucleofector T cell solution with 5-10 µg of NotI-linearized plasmid (New England Biolabs). Parasites were grown for 6 days in culture medium and selected for resistance using 5.0µg/ml Blasticidin. T7 promoter driven guide RNA constructs were linearized and introduced using the same method except 2.5µg/ml phleomycin was used for selection. Expression of each introduced protein was confirmed by western blot (see below).

### Plasmid construction

The plasmid containing the dCas9 Bdf3 fusion sequence was constructed in two halves: the first half consisted of the 3X FLAG tag, La Nuclear Localization Signal (89), and the front half of the dCas9 coding region, and the second half consisted of the back half of the dCas9 coding region, the GS linker, and the Bdf3 coding region. Both halves were synthesized by Twist Biosciences. The front 3X Flag/La NLS/dCas9 coding region was amplified using Forward ggccaagcttatggactacaaggaccac and Reverse gcgcgggcccacgtagtacggg primers containing HindIII (New England Biolabs) and ApaI (New England Biolabs) restriction enzyme recognition sites. This section was digested with HindIII and ApaI and cloned into a pLEW100 vector at these same sites. The back dCas9/Glycine-Serine linker/Bdf3 coding region was cut with ApaI and BamHI (New England Biolabs) and cloned into the pLEW100/3X Flag/La NLS/dCas9 vector at the ApaI and BamHI sites within the vector.

### sgRNA construction

Complementary single-stranded oligos were engineered with BbsI overhangs. Oligos were combined at a concentration of 5µM each in TE buffer with a reaction volume of 50µL. The oligos were annealed in a thermocycler as follows: 93°C for 0:01 with a ramp time of 3 °C/s, 70 °C for 3:00 with a ramp time of 3 °C/s, and a hold at 4°C with a ramp time of 0.1 °C/s. pT7sgRNA (56) was a gift from David Horn (Addgene plasmid # 111820 ; http://n2t.net/addgene:111820 ; RRID:Addgene_111820). pT7sgRNA vector was linearized with BbsI-HF (New England Biolabs) and purified. 150 ng of linearized pT7sgRNA and annealed oligo at a concentration of 2 µM of were ligated together using quick ligase (New England Biolabs) in a 20µL reaction for 5 minutes at 25°C. Ligated plasmids were transformed into electrocompetent cells to screen for correct integration of the oligos. The oligo sequences used were: EP1-1 forward, agggtgggcgtgcattgaaaatag; EP1-1 reverse, aaacctattttcaatgcacgccca; EP1-3 forward, aggggctgttccgtgtctctgggt; EP1-3 reverse, aaacacccagagacacggaacagc; Neo1 forward, aggggatctggacgaagagcatca; Neo1 reverse, aaactgatgctcttcgtccagatc.

### Western Blotting

SM *EPI/GFP* dCas9-Bdf3, dCas9 alone, or Bdf3 alone parasites were treated with or without 1 µg/mL dox and 8 million parasites were harvested after 48h or 72h of treatment. Harvested parasites were resuspended in 2X Laemmli buffer and boiled at 95°C for 10 minutes. The samples were loaded onto a 4-20% gradient polyacrylamide gel and protein was transferred to the PVDF membrane. The membrane was blocked in 7 mL of 5% milk in Tris-Buffered Saline (TBS), then incubated overnight at 4°C in M2 mouse anti-FLAG (Sigma), rabbit anti-FLAG (Cell Signaling), or the control anti-tubulin (a kind gift from George Cross) in 5% milk in (TBS) at a 1:1000 dilution. After 3 washes in TBS with 0.1% Tween-20, the membrane was incubated at room temperature for 1 hr in goat anti-mouse or goat anti-rabbit fluorescently labeled antibody (Biorad) at a 1:10000 dilution in 5% milk in TBS. After all antibody incubations, the membrane was washed three times in TBS with 0.1% Tween-20 and a final wash in TBS alone. The membrane was imaged on a LI-COR Odyssey instrument.

### Flow cytometry

SM *EPI/GFP* dCas9-Bdf3 or dCas9 cells were prepared at a density of 15,000 cell/mL and treated with or without 1 μg/mL dox in 2 mL of culture medium. Parasites were diluted in fresh media with dox as needed to maintain them under 1 million/ml. After two and three days of growth, 1 mL of culture was harvested and analyzed by flow cytometry for *EP1/GFP* expression. All samples were analyzed with a Novocyte 2000R from Acea Biosciences.

### Analysis of parasite growth

Parasites were prepared at a density of 100,000 cells/mL and grown in 96 well plates with a starting volume of 200µL HMI-9 medium for 24h with or without 1 µg/mL dox. After 24 hours of growth, 20 µL of culture was analyzed by flow cytometry and the number of parasites in the live gate was calculated. Cultures were diluted back down to 100,000 cell/mL using the number of cells determined by the live gate. This procedure occured daily for a total of 5 days.

### Quantitative PCR

RNA was isolated from parasites using the Direct-zol RNA MiniPrep kit from Genesee, which includes DNAse treatment. cDNA was generated using the SuperScript IV VILO Master Mix (Fisher Scientific) according to the manufacturer’s instructions starting with 1µg of RNA. cDNA was amplified with primers combined with 2X Sybr green master mix (Life Technologies) and analyzed on an Eppendorf Realplex2 instrument. Primers used were as follows: Tb427.10.10260 EP1, tctgctcgctattcttctgttc, cctttgcctcccttagtaagac, Tb927.6.510 GPEET agtcggctagcaacgttatc, ttctggtccggtctcttct, Tb927.9.11600, GIM5B, ttgcgaggatgggtgatg, gggtttggagagggaagttaat, Tb927.10.2010, HK1, gtcagcacttactcccatcaa, acgacgcatcgtcaatatcc, Tb927.10.5620, ALD, gtctgaagctgttgttcgtttc, cacctcaggctccacaatag, Tb927.10.10220, PAG2, aggagatacgaggaatgagaca, tcttcaaacgcccggtaag, Tb927.10.14140 PYK1, gagaaggttggcacaaagga, tcacaccgtcgtcaacataaa, GFP, ctacaacagccacaaggtctat, ggtgttctgctggtagtg, Tb927.10.9400, SF1, ggtatggttcatcaggagttgg, cgttagcactggtatccttcag.

### Accession numbers

EP1, Tb927.10.10260

PAG2, Tb927.10.10220

GPEET, Tb927.6.510

HK1, Tb927.10.2010

GIM5B, Tb927.9.11600

ALD, Tb927.10.5620

PYK1, Tb927.10.14140

SF1, Tb927.10.9400

BDF3, Tb927.11.10070

## Acknowledgements.

We would like to acknowledge Monica Mugnier, who provided excellent advice on adapting existing CRISPR-Cas9 systems to *T. brucei*. D.S. is supported by a NSF CAREER grant (2041395), which also supported the undergraduate students who worked on the project.

## Notes

### Competing Interest Statement

The authors have declared no competing interest.

### Summary of Updates

The previous version did not include supplemental data. This version includes supplemental data.

