## Supplementary material for "The bromodomain protein Bdf3 is sufficient to activate expression of a procyclin gene in bloodstream stage African trypanosomes": S1 Figure and Legends

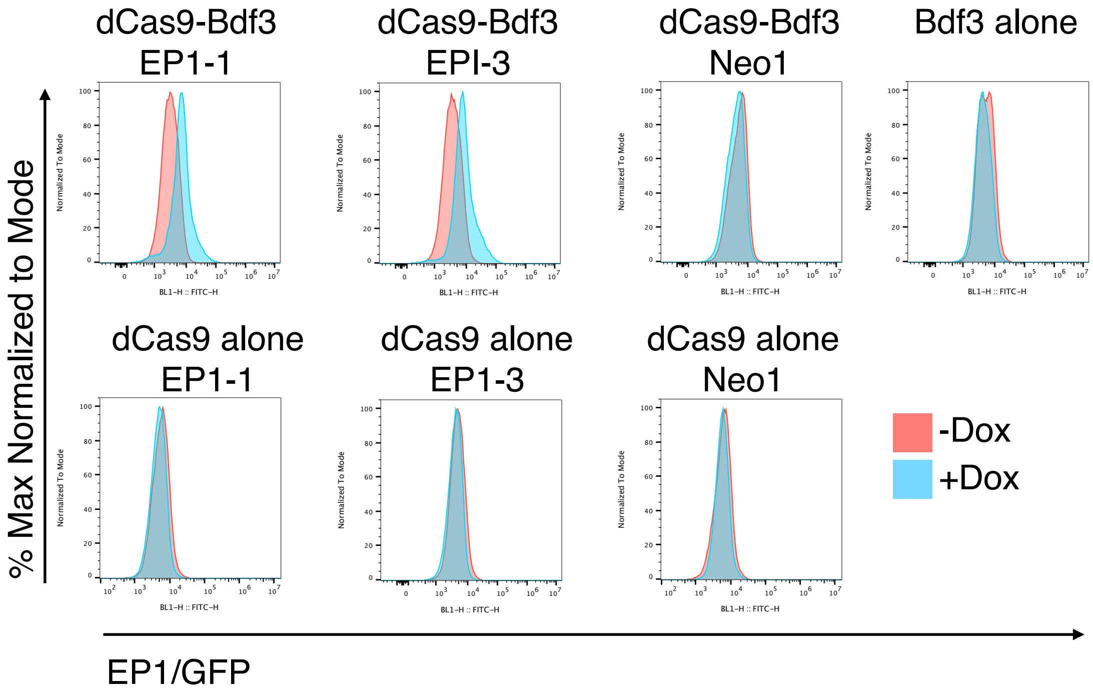


S1 Figure. Representative histograms generated from flow cyometric analysis of parasites transfected with each indicated inducible expression plasmid and guide RNA. Parasites were treated for 3 days with 1$\mu$g/ml doxycyclin to induce expression of the indicated protein and the guide RNA. -Dox parasites represent negative controls.

S1 Table. p-values generated from ANOVA statistical test followed by Tukey HSD post-hoc test on flow cytometric data from parasites transfected with inducible dCas9 constructs and guide RNAs treated with 1$\mu$g/ml doxycyclin for 48h.

S2 Table. p-values generated from ANOVA statistical test followed by Tukey HSD post-hoc test on flow cytometric data from parasites transfected with inducible dCas9 constructs and guide RNAs treated with 1$\mu$g/ml doxycyclin for 72h.
